## Supplementary for "Spatiotemporal variations in retrovirus-host interactions among Darwin’s finches"

**Supplementary Table 1 | ERV locus density by chromosome.**

| chromosome | number of loci | chromosome size | ERV loci per MB |
|---|---|---|---|
| chr2 | 3149 | 152240728 | 20,68 |
| chr1 | 2732 | 113967701 | 23,97 |
| chr3 | 2417 | 113435954 | 21,31 |
| chrZ | 1888 | 73580794 | 25,66 |
| chr1A | 1708 | 72438334 | 23,58 |
| chr4 | 1567 | 71061686 | 22,05 |
| chr5 | 1389 | 62010874 | 22,40 |
| chr7 | 699 | 38189422 | 18,30 |
| chr6 | 751 | 35412503 | 21,21 |
| chr8 | 644 | 30584378 | 21,06 |
| chr9 | 560 | 25423666 | 22,03 |
| chr11 | 487 | 20882383 | 23,32 |
| chr12 | 417 | 20836500 | 20,01 |
| chr10 | 249 | 20345023 | 12,24 |
| chr4A | 404 | 19694112 | 20,51 |
| chr13 | 314 | 18340482 | 17,12 |
| chr14 | 327 | 16472583 | 19,85 |
| chr20 | 531 | 14936690 | 35,55 |
| chr15 | 439 | 14104672 | 31,12 |
| chr18 | 383 | 12083600 | 31,70 |
| chr17 | 519 | 11493945 | 45,15 |
| chr19 | 388 | 11160396 | 34,77 |
| chr21 | 239 | 7910879 | 30,21 |
| chr24 | 442 | 7709400 | 57,33 |
| chr23 | 388 | 7058734 | 54,97 |
| chr26 | 365 | 6857747 | 53,22 |
| chr28 | 343 | 5889000 | 58,24 |
| chr27 | 265 | 5560827 | 47,65 |
| chr22 | 344 | 4946000 | 69,55 |
| chr25 | 78 | 3215879 | 24,25 |

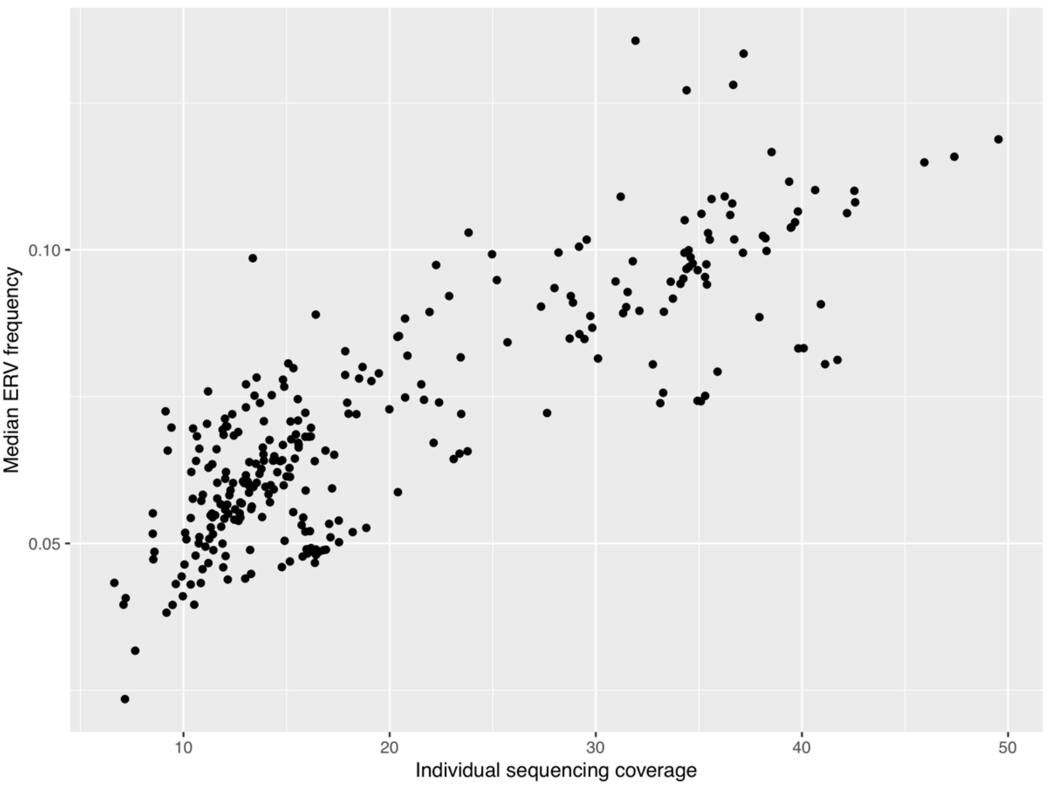


**Supplementary Fig. 1 | Relationship between ERV detection by RetroSeq and mapped coverage.** Plot of mean whole-genome sequencing mapped coverage (x) and the number of detected RetroSeq ERV insertion loci called per individual.

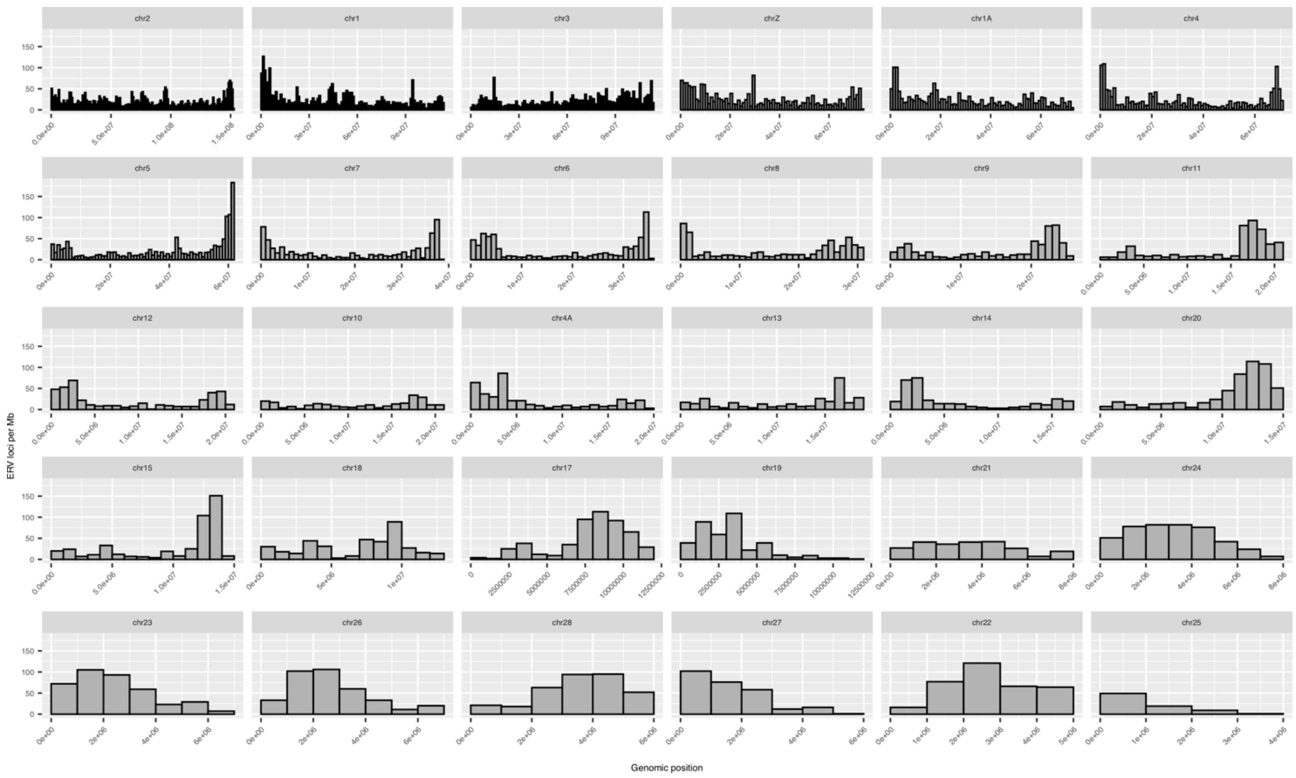


**Supplementary Fig. 2 | ERV genome density by chromosomes.** Non-uniform overall ERV distribution density histograms showing differences between and along chromosomes.

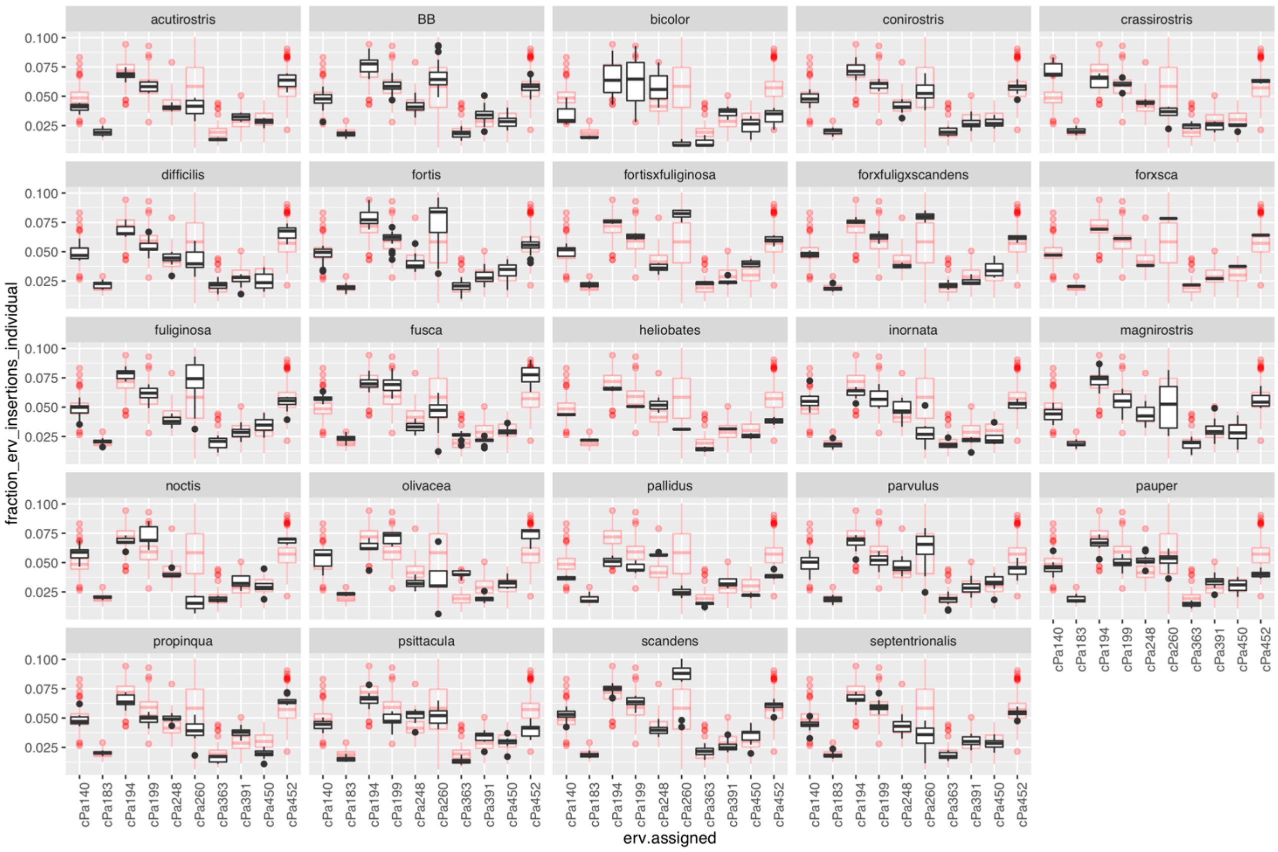


**Supplementary Fig. 3 | Relative abundance of common ERVs.** The 10 most frequent ERVs showed significant variation in abundance both within and between species. The relative fraction of insertions of an ERV within an individual was plotted as a data point in the box plots. Red data points are for all finch samples, and black is the subset corresponding to only the species in the labeled window. Variation in relative ERV abundance among all samples (e.g. cPa260), could be attributed to either within species variation (e.g. *G. fortis* and *G. magnirostris*), between species variation (e.g. *L. noctis* and *G. scandens*), or both.

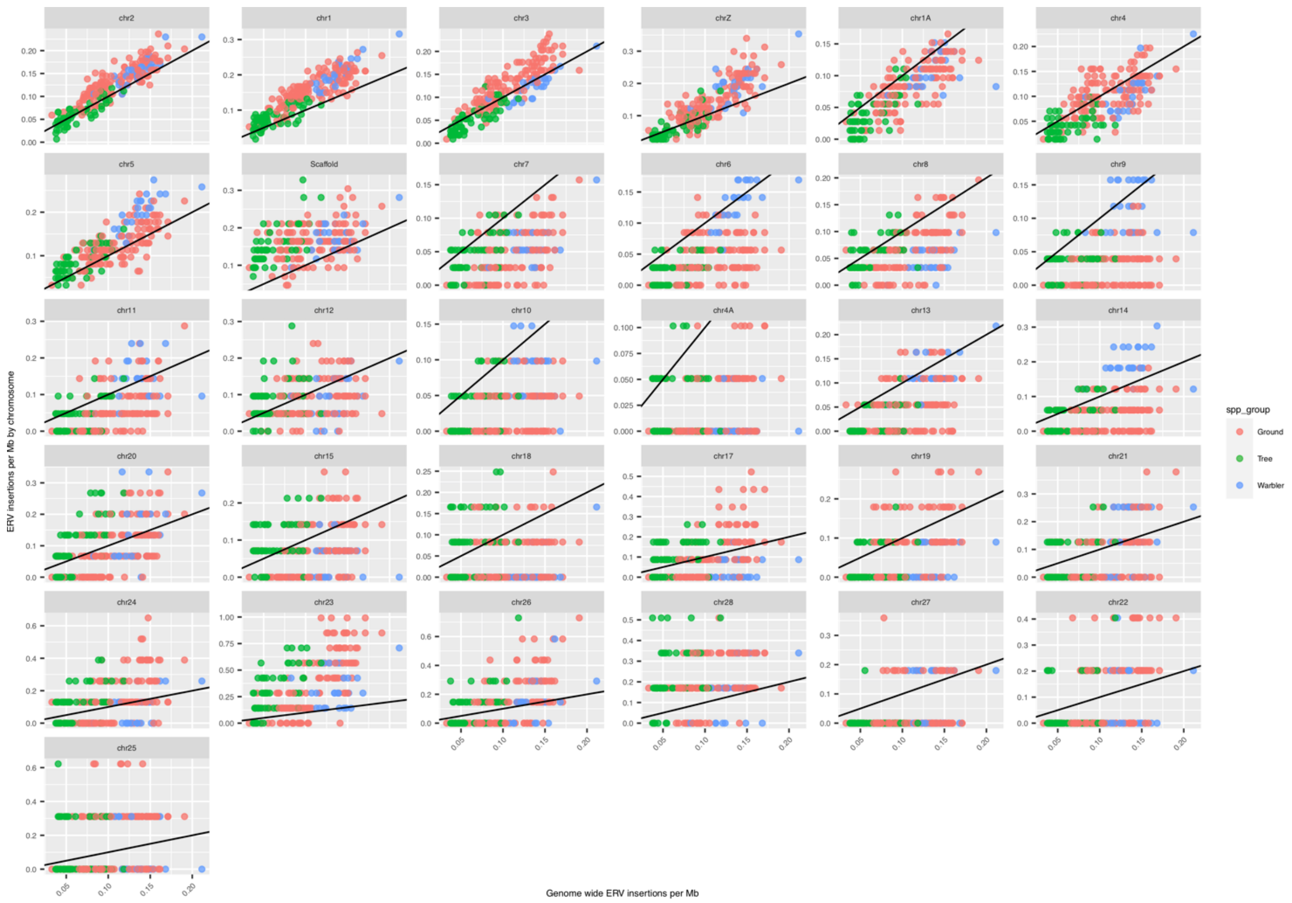


**Supplementary Fig. 4 | Density of cPa452 by chromosome for warbler-, tree- and ground finches.** Density on each chromosome of *Beta-like 1* ERV cPa452 compared to overall density for each individual sample of ground finch (red color), tree finch (green color), and warbler finch (blue color). Points above the line indicate that cPa452 occurs with a higher density on the chromosome indicated than in the overall genome, where density is measured in ERV insertions per Mb.

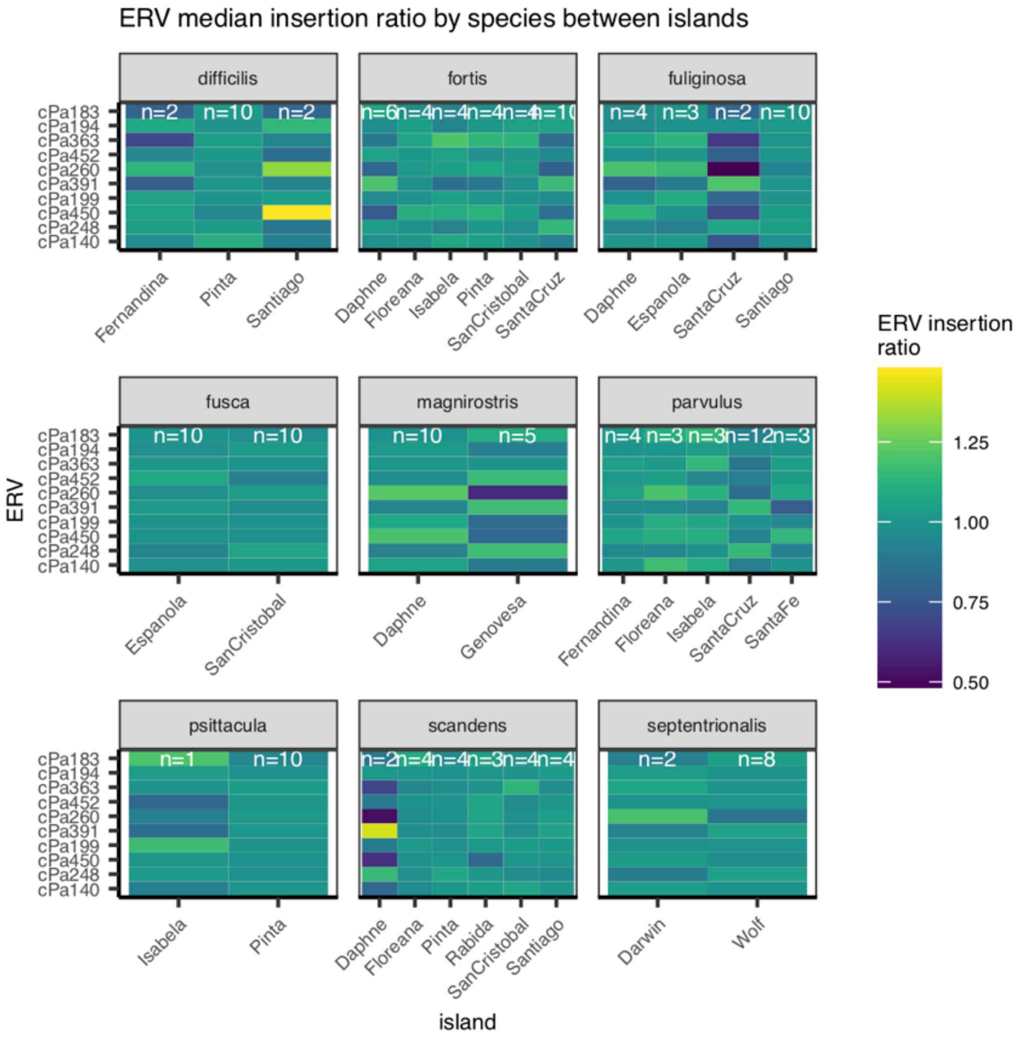


**Supplementary Fig. 5 | Contrasting ERV landscapes across finches and islands.** Relative abundance of the 10 most frequent ERVs showed variation between island populations of the same species. Modified MIR normalized ERV abundance values both vertically across ERVs and horizontally across islands. A contrasting color between island populations for a given ERV indicates a difference in relative abundance between populations, either higher (lighter color), or lower (darker color). Variation in MIR increased with small sample size (e.g. *G. scandens* from Daphne island), however some contrasts are more likely to represent large actual differences in ERV abundance between island populations (e.g. cPa260 in *G. magnirostris*).

**Supplementary Data 1 | ERV FASTA sequences.**

**Supplementary Data 2 | ERV nexus tree file.**
